# Re-investigating the correctness of decoy-based false discovery rate control in proteomics tandem mass spectrometry

**DOI:** 10.1101/2023.06.21.546013

**Authors:** Jack Freestone, William Stafford Noble, Uri Keich

## Abstract

Traditional database search methods for the analysis of bottom-up proteomics tandem mass spectrometry (MS/MS) data are limited in their ability to detect peptides with post-translational modifications (PTMs). Recently, “open modification” database search strategies, in which the requirement that the mass of the database peptide closely matches the observed precursor mass is relaxed, have become popular as a way to find a wider variety of types of PTMs. Indeed, in one study, Kong *et al*. reported that the open modification search tool MSFragger can achieve higher statistical power to detect peptides than a traditional “narrow window” database search. At the same time, Kong *et al*. reported that their empirical results suggest a problem with false discovery (FDR) control in the narrow window setting. We investigated these claims empirically and, in the process, uncovered a potential problem with FDR control in the machine learning post-processors Percolator and PeptideProphet. However, we also found that, after accounting for chimeric spectra as well as for the inherent difference in the number of candidates in open and narrow searches, the data does not provide sufficient evidence that FDR control in proteomics MS/MS database search is problematic.

## 1 Introduction

The proteomics search engine MSFragger was developed to help detect post-translationally modified peptides [12]. Such peptides pose a challenge because using a traditional “narrow window” database search to detect them would involve an infeasible, exponentially large search space. Instead, MSFragger efficiently employs “open modification” search, in which each observed spectrum is compared against peptides whose masses differ—often by hundreds of Daltons—from the observed precursor mass associated with the spectrum [1]. Kong *et al*. demonstrated that MSFragger can achieve higher statistical power to detect peptides than a traditional “narrow window” database search [12]. In the process, Kong *et al*. also claimed to have uncovered a liberal bias in the FDR control provided by standard procedures for MS/MS database search.

We investigated these two observations empirically and reached several related conclusions. First, our analysis shows that open searches on their own often produce fewer discoveries than narrow searches applied to the same data. Only when coupled with a machine learning post-processor such as Percolator [9] or PeptideProphet [10] do open searches become typically better than narrow window searches. We hypothesized that this discrepancy arises in part due to a problem with false discovery rate (FDR) control. Accordingly, using entrapment experiments [8] we show that in practice both Percolator and PeptideProphet can apparently fail to control the FDR. Hence, the apparent improvement in power attributed to open modification searching must be taken with a grain of salt.

Motivated by these observations, we recently developed an alternative post-processing approach that allows us to control the FDR in a paired set of narrow and open search discoveries. Notably, our new method, Combining Open and Narrow searches with Group-wise Analysis (CONGA), can also detect and better utilize chimeric spectra that can identify multiple peptides [6]. With CONGA’s help we revisit Kong *et al*.’s claim that open search uncovers a problem with decoy-based FDR control in narrow search [12]. Specifically, we find that after accounting for chimeric spectra, as well as for the inherent difference in the number of candidates in open and narrow searches, the data does not provide sufficient evidence that the FDR in the narrow search is uncontrolled.

## 2 Methods

### 2.1 Datasets

#### 2.1.1 Entrapment run searches

Entrapment runs for Percolator and PeptideProphet were performed using data taken from the standard protein mix database, ISB18 [11]. We used the nine `.ms2` spectrum files taken from [13] which were originally sourced from the Mix 7 spectrum files downloaded from https://regis-web.systemsbiology.net/PublicDatasets/18_Mix/Mix_7/ORBITRAP/ (this excluded the tenth smallest spectrum file). These `.ms2` spectrum files were subsequently used for Tide. For compatibility with MSFragger, we directly downloaded the same nine spectrum files in ThermoRaw format and converted them to a single combined, as well as separate, `.mzML` files using MSConvert 3.0.22314 with the vendor peak-picking filter using the default settings.

The in-sample database comprised of the extended protein database containing the 18 peptides and 30 additional hitchhiker peptides downloaded from https://regis-web.systemsbiology. net/PublicDatasets/database. We used the castor plant proteome as the entrapment database as in [13]. Tide-index was used to digest the in-sample and entrapment database using trypsin, for Tide, and strict-tryspin (specified with `trypsin/p`), for MSFragger. For each digest rule, four random subsets of the in-sample peptides of varying size, plus the entrapment peptides were used to create a combined target peptide database for which 20 randomly shuffled decoy indices were created, again using tide-index. Param-medic [16] was used to determine any variable modifications, though none were detected. Any entrapment peptide that was identical (up to any leucine/isoleucine substitution) to an ISB18 peptide was removed from the entrapment database. Table 1 lists the resulting number of in-sample and entrapment peptides in each combined target peptide database, along with their in-sample-to-entrapment ratio.

**Table 1:** The number of in-sample target peptides and entrapment-peptides for each combined target peptide database used in the entrapment runs. The table reports the number of in-sample target peptides and entrapment target peptides for each of the four combined target databases, for each digest rule. The first column reports the proportion of the total in-sample target peptides used in the combined target database, the second is the number of these in-sample peptides used, the third column is the number of entrapment peptides, and the last column is the ratio of in-sample peptides to entrapment peptides.

| Trypsin for Tide |  |  |  |
| --- | --- | --- | --- |
| In-sample proportion | In-sample peptides | Entrapment peptides | In-sample:entrapment |
| 100% | 898 | 571298 | 1:636 |
| 75% | 674 | 571298 | 1:848 |
| 50% | 449 | 571298 | 1:1272 |
| 25% | 225 | 571298 | 1:2539 |
| Strict-trypsin for MSFragger |  |  |  |
| In-sample proportion | In-sample peptides | Entrapment peptides | In-sample:entrapment |
| 100% | 919 | 584350 | 1:636 |
| 75% | 690 | 584350 | 1:847 |
| 50% | 460 | 584350 | 1:1270 |
| 25% | 280 | 584350 | 1:2087 |

Tide-search was used to search the nine `.ms2` spectrum files against each of the combined target and decoy databases prepared using trypsin with the following settings: for narrow searches we used `--auto-precursor-window warn --auto-mz-bin-width warn --use-tailorcalibration T --concat T`, and for open searches we used `--auto-mz-bin-width warn --precursor-window-type mass --precursor-window 100 --top-match 5 --concat T -use -tailor-calibration T`. MSFragger v3.3 was used to search the combined `.mzML` and the separate `.mzML` spectrum files against each of the combined target and decoy databases prepared using strict-trypsin. Because we used tide-index to prepare these databases, we need to use the following settings to ensure MSFragger has the same digestion parameters. For narrow searches we used `allowed_missed_cleavage = 0, digest_min_length = 6, digest_max_length = 50, digest_mass_range = 200.0 7200.0, allowed_missed_cleavage = 0` with all other options set to default, and for open searches we used `localize_delta_mass = 0, allowed_missed_cleavage = 0, digest_min_length = 6, digest_max_length = 50, digest_mass_range = 200.0 7200.0, allowed_missed_cleavage = 0, output_report_topN = 5`, with all other option set to default. No variable modifications were set, so CONGA’s analysis was done at the peptide level. Note we used MSFragger v3.3 at the time, as the subsequent version at the time (v3.4) did not allow for `output_report_topN > 1` for DDA data. We used Tide in Crux v4.1.decd99ff.

#### 2.1.2 PRIDE-20 searches

We downloaded 20 high-resolution spectrum files from the Proteomics Identifications Database, PRIDE [15]. Seven of these spectrum files were taken from [7] and were originally obtained by randomly selecting a spectrum file from randomly selected PRIDE projects (submitted no earlier than 2018). The remaining 13 spectrum files were similarly obtained by randomly selecting a single spectrum file from a randomly selected PRIDE project (submitted no earlier than 2019). The sampling was constrained to generate a collection of 10 spectrum files that had modifications and 10 spectrum files for which no modifications were detected, as determined in both cases by Parammedic [16]. The protein FASTA database files were also downloaded from the associated PRIDE projects or in the case of human data, the UniProt database UP000005640 was used (downloaded 9/11/2021). Table 2 reports the list of the 20 spectrum files and PRIDE projects used.

**Table 2:**
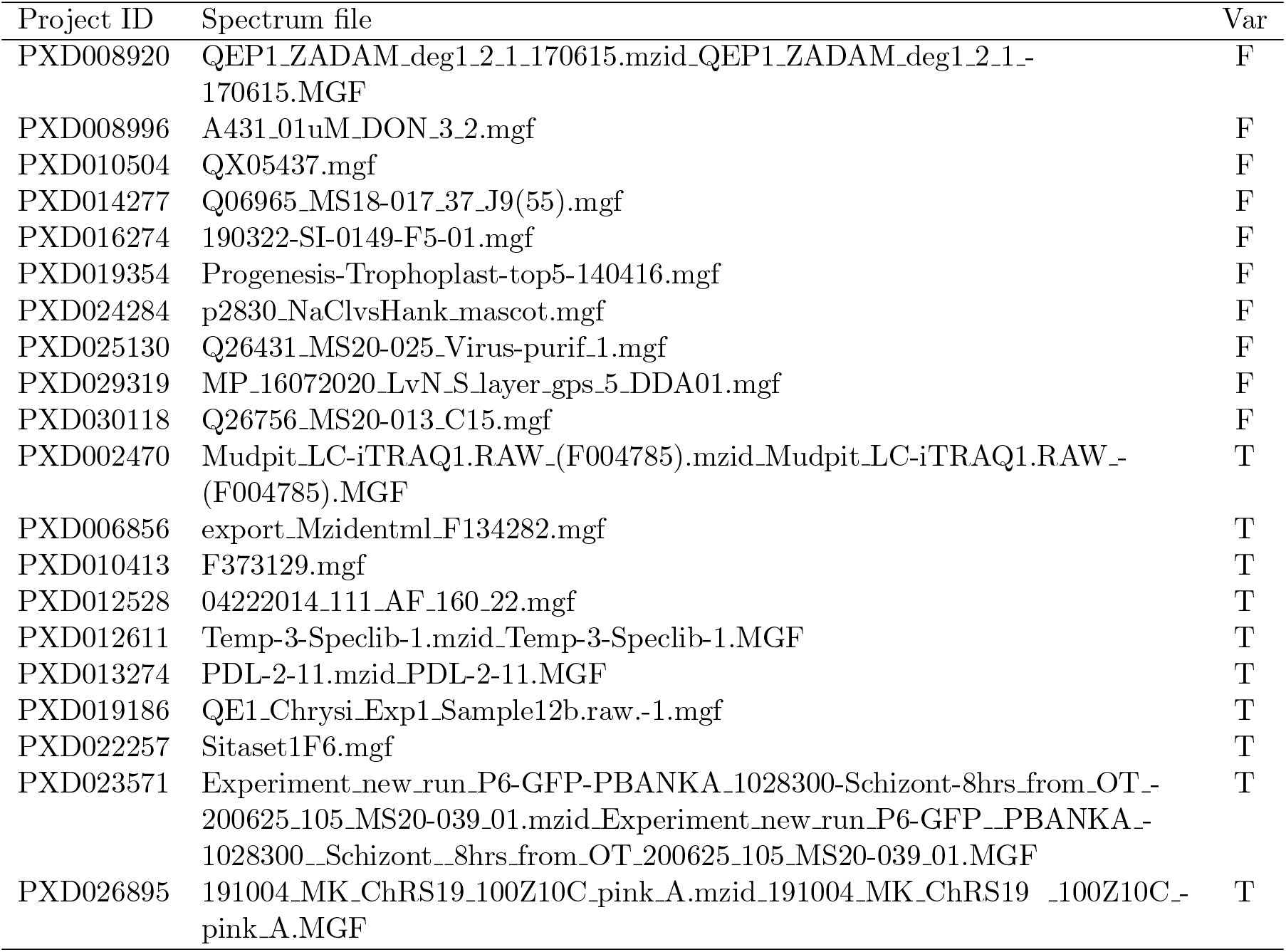
The PRIDE-20 data set. The list of 20 spectrum files used in the PRIDE-20 data set and their associated project IDs. The “Var” column indicates whether the data includes variable modifications.

For each of the PRIDE-20 data sets, we used tide-index 20 times to create 20 different target-decoy peptide databases using `--auto-modifications T` to call Param-medic and apply the variable modifications. For narrow searches using Tide, we used the following options `--auto-precursor-window warn --auto-mz-bin-width warn --use-tailor-calibration T --concat T` and for open searches using Tide we used `--auto-mz-bin-width warn --precursorwindow-type mass --precursor-window 100 --use-tailor-calibration T --concat T`.

For searches using Comet, we used the builtin version within Crux. For narrow searches we used the options `--decoy search 1 –auto_peptide mass tolerance warn --allowed_missed_cleavage 0 –auto_fragment bin tol warn –auto_modifications T --peptide mass units 2` and for open searches we used `--decoy search 1 –peptide_mass tolerance 100 --allowed_missed_cleavage 0 –auto_fragment_bin_tol warn –auto_modifications T –peptide_mass_units 0`. Comet digests and produces decoy peptides by reversing each target sequence. Hence, only a single target-decoy database was used for each of the PRIDE-20 data sets. PSMs matched to peptides that were identified as both a target and decoy were deleted. In the case of E-values, lower scores indicate a better PSM and so we take the negative of these scores when applying TDC.

Lastly for searches using MSFragger we converted each `.mgf` file into `.mzmL` files using MSConvert with default settings. We constructed a decoy protein database by taking each target protein and reversing the position of the amino acids that lie between *K, R* or the *N* - and *C*-terminal, while keeping the position of *K* and *R* fixed. Importantly, the digestion of the target and decoy proteins allow us to pair each resulting target peptide with a corresponding decoy peptide. This is not achievable if we reverse the entire target protein sequence, which is the behaviour of many softwares. For narrow searches, we used the following options `allowed_missed_cleavage = 0, use_topN peaks = 100` and for open searches we used `output_report_topN = 5, allowed_missed_cleavage = 0, use_topN_peaks = 100`. Variable modifications were set according to those detected by Parammedic using tide-index. All other options were set to the default settings. Similar to Comet, PSMs matched to peptides that were identified as both a target and decoy were deleted.

#### 2.1.3 HEK293 searches

We downloaded the 24 HEK293 spectrum files from https://ftp.pride.ebi.ac.uk/pride/ data/archive/2015/06/PXD001468 which were converted to `.mzML` format using MSConvert with the vendor peak-picking filter using the default settings. We used the UniProt database UP000005640 (downloaded 18/05/2022). As discussed in Section 2.1.2, we prepared a combined target-decoy protein database by taking each target protein and reversing the position of the amino acids that lie between *K, R* or the *N* - and *C*-terminal, while keeping the position of *K* and *R* fixed. We ran narrow MSFragger searches for each of the 24 spectrum files using the settings `precursor_mass_lower = -100, precursor_mass_upper = 100, allowed missed cleavage = 1, use_topN_peaks = 100` and open searches using the settings `precursor_mass_lower = -500, precursor_mass_upper = 500, precursor_mass_units = 0, localize_delta_mass = 0, allowed_missed_cleavage = 1, output_report_topN = 5`. No variable modifications were considered, and all other options were set to default. These settings were used to follow as best we reasonably could the setup used by [12].

#### 2.1.4 Target-decoy competition

For the TDC procedure, we employ the double competition protocol, “PSM-and-peptide” as originally described in [13], and summarized for convenience below as well in pseudocode in Supplementary Algorithm 1.^1^ For the PSM-level competition, we determine the best peptide-spectrum match to the target database and decoy database separately, and then record the best scoring peptide-spectrum match out of the two. This is achieved implicitly by taking the top 1 PSM for each spectrum when searching against the concatenated target-decoy database. For the subsequent peptide-level competition, we define the score of each peptide as the maximal PSM score associated to that peptide. Then for each target-decoy pair, we take the peptide with highest peptide score. The resulting winning peptides are then ordered according to their score from largest to smallest. Denoting the sorted winning scores as *W*_1_ *≥ … ≥ W*_*n*_, we determine the largest index *K* for which the estimated FDR is *≤* the threshold *α*, and report all target winning peptides up to and including *K*. More specifically, denoting *L*_*i*_ = 1 as a target (winning) peptide and *L*_*i*_ = *−*1 as a decoy peptide, then all target peptides with index *i ≤ K* are reported, where

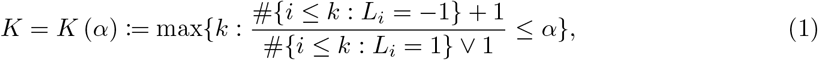

and *x ⋁ y* := max*{x, y}*.

#### 2.1.5 Percolator settings

For each of the Tide searches where Percolator was subsequently used, we constructed `.pin` files by either specifying `--pin-output T` during Tide search or by using the `make-pin` utility command in Crux. When creating `.pin` files from an open search we used the default number of top 5 PSMs which allows for generating the feature `deltLCn` (the difference between the XCorr score of the top 1 PSM and the last ranked PSM). We then kept only the top 1 PSM for each spectrum to overcome the problem of neighbors. For narrow searches, we directly used the top 1 PSM. Lastly, to each PSM in the `.pin` file, we set the `enzInt` feature (the number of internal enzymatic sites) to zero as explained in [3]. For MSFragger-searches, we specified `output_format = tsv_pepxml_pin` and subsequently filtered for the top 1 PSM as above.

For each of the .pin files, we used the following options in the crux percolator command `--only-psms T, --tdc F`. This reports only the PSMs and does not apply the TDC procedure within Percolator since it does not use the double competition protocol, “PSM-and-peptide”. Instead, we apply our TDC procedure as described in the previous Section 2.1.4 to the resulting list of PSMs, using the learned percolator score as the PSM score.

#### 2.1.6 PeptideProphet

We used the Philosopher software to apply PeptideProphet to MSFragger searches of the entrapment runs. We followed Philosopher’s workflow, using the following commands. To initialize the workspace, we used: `workspace --init`. To store the database in the workspace we used `database ---prefix decoy_ --nodecoys --custom fasta_file` where the `fasta_file` is the combined target-decoy peptide database containing a random subset of the in-sample peptides and the entrapment peptides (details given in Section 2.1.1). Then to run PeptideProphet we used `peptideprophet --database prepared_fasta_file --expectscore --decoyprobs --decoy decoy_ --nonparam --mass width 1000 search file` where `prepared fasta_file` is the database generated in the previous step and `search file` corresponds to the MSFragger search files used in the entrapment runs (details given in Section 2.1.1). The FDR analysis was conducted in one of two ways: (1) using TDC with PeptideProphet’s linear discriminant score (denoted as `fval` in the output) in lieu of the PSM score, and (2) using Philosopher’s validation. In the latter case, we used the command `filter --tag decoy --pep x --psm x --pepxml pp_file` where `pp_file` is the output of PeptideProphet in the previous step, and *x* is the threshold ranging from 0.01 to 0.1.

## 3 Results

### 3.1 In the absence of a post-processor, open search is not more powerful than narrow search

Motivated by the increasing popularity of open modification searching, we set out to compare the statistical power of open and narrow window searches to detect peptides from MS/MS data. We reasoned that, although open search allows for the discovery of unexpected PTMs, it also necessarily considers many more candidate peptides for each observed spectrum. This large number of candidates increases the risk that a correct match—that is, a match to the peptide that truly generated the observed spectrum—may be out-scored by an incorrect match with a randomly high score. The empirical question boils down to whether the increased rate of discoveries of peptides with unexpected PTMs outweighs the loss of power due to the larger number of candidate peptides in the open search.

To investigate this question, we searched a collection of publicly available data using three different search engines. For the data, we used 20 high-resolution spectrum files, each from a distinct project from the Proteomics Identifications Database (PRIDE) [15]. These data sets were selected essentially at random, subject to the constraint that 10 of the spectrum files required variable modifications, whereas the other 10 did not. We then applied three search engines—Tide [4], Comet [5], and MSFragger [12]—to each dataset, in both narrow and open mode. Each search employed a concatenated target-decoy database, where Tide used shuffled peptides as decoys, and MSFragger and Comet used reversed peptides as decoys. Furthermore, to obtain more accurate estimates, we repeated each Tide search 20 times, using 20 independently shuffled decoy databases, each concatenated with a fixed set of targets (see Section 2.1.2 for details).

For each search engine the reported list of discovered peptides was generated using our recently described double competition protocol, “PSM-and-peptide” [13]. In this protocol, the first competition is at the PSM level, where each spectrum is searched against the concatenated target-decoy database and only the top matching PSM is kept. This is followed by a second, peptide-level competition, where each target peptide is associated with its paired decoy, and from each pair only the peptide with the highest PSM score is kept. TDC is then applied to the list of winning peptides sorted according to their maximal PSM scores (Section 2.1.4).

The results of this experiment show that we often obtain fewer discoveries using open search compared to narrow search (Figure 1). For all three search engines, at an FDR threshold of *α* = 1% we overall obtain more discoveries with narrow search than with an open one (i.e., both the mean and the median ratios are *>* 1). While for higher FDR thresholds the picture is murkier, it is still clear that a narrow search can often deliver substantially more discoveries than an open one and that the magnitude of the difference varies by dataset and search engine used. Strikingly, for MSFragger the mean of the ratio of the number of narrow-to-open search discoveries is much higher than the median and even higher than the 75th percentile, indicating that for a few of the PRIDE-20 datasets there were significantly more narrow than open discoveries.

**Figure 1.**
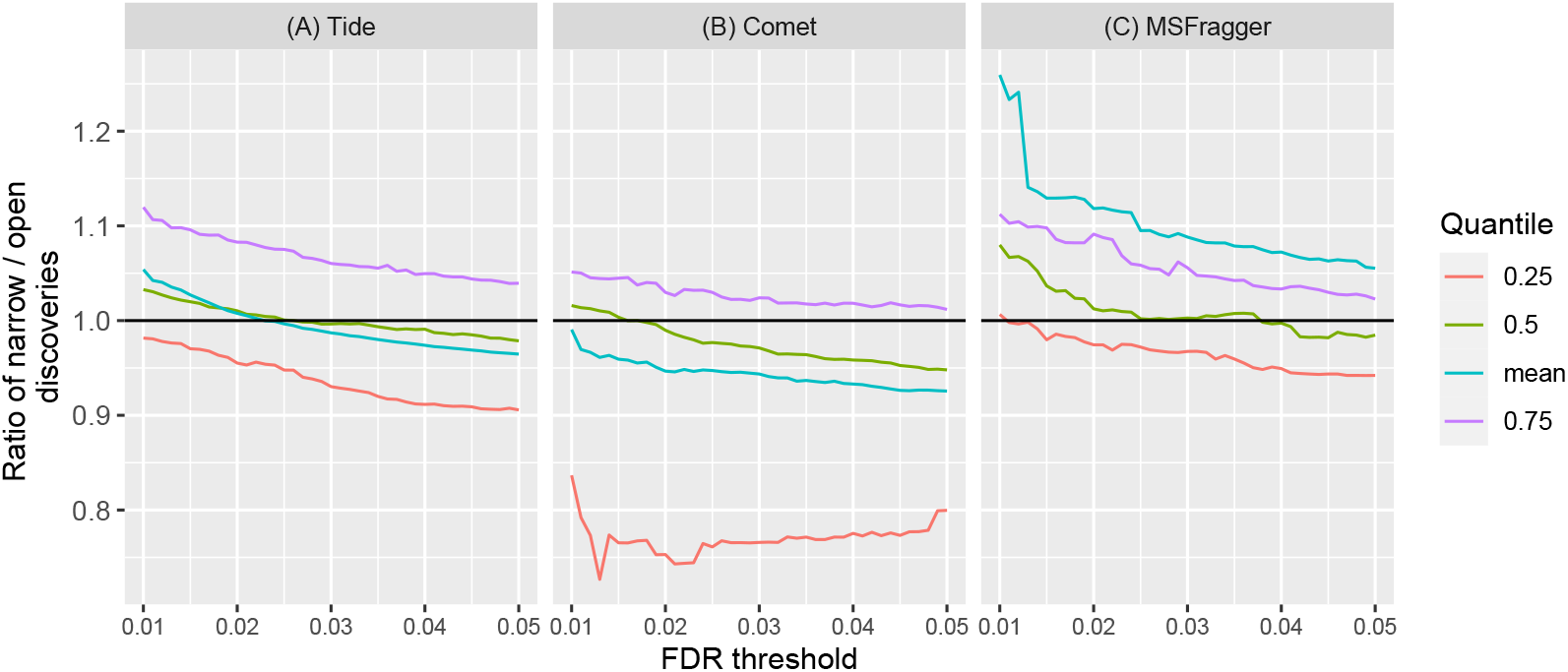
Quartiles of narrow-to-open search discoveries in real data. The mean and quartiles over the PRIDE-20 datasets of the ratio of narrow-to-open search discovered peptides using (A) Tide, (B) Comet, and (C) MSFragger, plotted as a function of FDR threshold. In (A), for each of the 20 datasets we average the discovery ratios over 20 randomized decoys generated specifically for that dataset, whereas in (B–C), we use a single reversed decoy database for each dataset. In panel (C) the mean is much higher than the median due to several outlier datasets.

### 3.2 Post-processors such as Percolator and PeptideProphet can struggle with FDR control

The results in Figure 1 are in apparent disagreement with results from Kong *et al*. [12], who reported greater power to detect peptides from open compared to narrow searches. One potential source of this discrepancy is that we carried out FDR control directly on the results of the search engine, whereas Kong *et al*. relied on FDR estimates from a machine learning post-processor. Two such post-processors, Percolator [9] and PeptideProphet [10, 14], are widely used to re-rank PSMs produced by a database search and carry out FDR control. We therefore sought to empirically assess the FDR estimates provided by these tools.

To do so, we ran a set of entrapment experiments, in which spectra that were generated in a controlled experiment from a known set of peptides are searched against a database containing those peptides plus a set of “entrapment sequences” [8]. In our case, we use spectra from a previously published analysis of an 18-protein mixture (ISB18) [11]. Furthermore, to simulate datasets with a varying degree of signal-to-noise ratio we constructed four different combined target databases using an increasingly smaller portion of the complete in-sample database. Specifically, we combined a fixed set of entrapment sequences with a randomly sampled proportion of the in-sample ISB18 database: 100%, 75%, 50%, and 25%. The ratio of entrapment to ISB18 peptides in those four target databases varied from 636:1 to 2539:1, ensuring that in all cases the vast majority of the false discoveries involve entrapment peptides. This setup allows us to reliably estimate the false discovery proportion (FDP) among the set of discovered peptides. Averaging the estimated FDP over 20 randomly generated decoy databases per entrapment target database (where each decoy database contains shuffled sequences of all the target sequences: ISB18 and entrapment ones), we obtain an estimated FDR that we can compare with the selected FDR threshold. If the FDR control procedure is valid, then this empirical FDR should not significantly exceed the threshold. We used both Tide and MSFragger to search each of 20 concatenated databases in both open and narrow modes. Percolator (both search engines) and PeptideProphet (MSFragger only) were then applied to rescore the PSMs to which we applied peptide-level TDC and estimated the FDR as above (see Section 2.1.1 for details). Both post-processors yield estimated FDRs that are consistently greater than the specified FDR threshold, across a range of thresholds from 0.01 to 0.1, and in both open and narrow searches (Figure 2A–C). Furthermore, as the number of in-sample peptides decreases, the estimated FDR is driven further up for most FDR threshold values. Note that because the ISB18 experimental spectra originated from very few proteins, for lower FDR thresholds and increasingly smaller in-sample portion we often obtain no discoveries. Thus, the larger FDR violations are observed at higher FDR thresholds.

**Figure 2.**
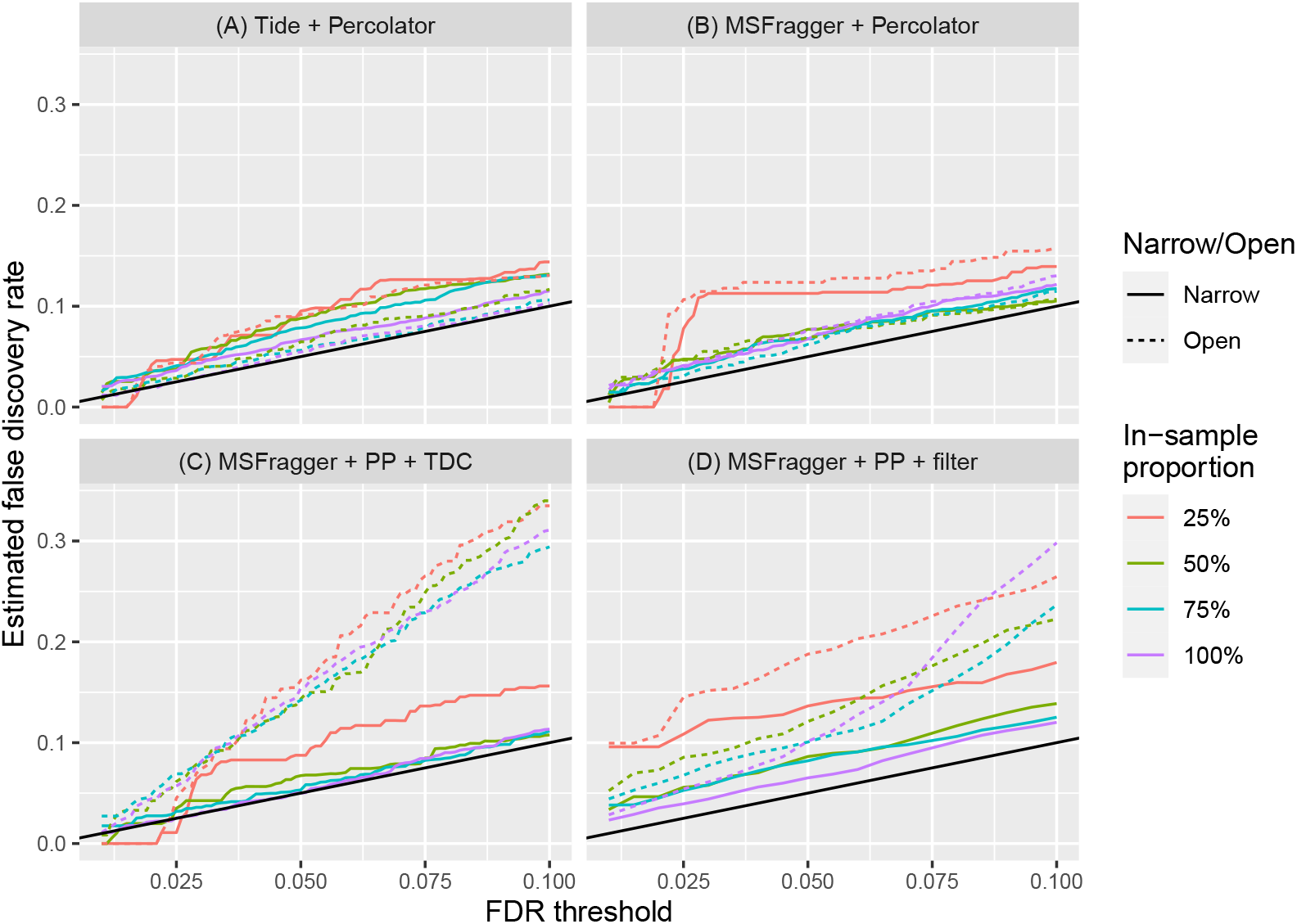
Percolator and PeptideProphet empirical FDR. The empirical FDR as estimated from our entrapment experiments on the ISB18 dataset using (A) Percolator with Tide, (B) Percolator with MSFragger, (C) PeptideProphet’s discriminant score with MSFragger, and (D) PeptideProphet with Philosopher’s default filter function with MSFragger. The FDP is estimated at a range of FDR thresholds ([0.01, 0.1]) and its average over 20 randomly generated decoys is the empirical FDR. Each curve represents the empirical FDR using a target database constructed with the specified proportion of the in-sample database, in either narrow-or open-search mode. The estimated FDRs mostly exceed the corresponding threshold (solid black line), indicating that the FDR is not properly controlled.

In addition, we also examined the default FDR analysis of the Philosopher workflow [2]. The latter relies on PeptideProphet’s posterior error probabilities to estimate and control the FDR (Section 2.1.6). We found that, particularly for lower FDR thresholds, Philosopher makes things even worse (Figure 2D); for example, the (worst-case) FDR at 1% changes from 2.72% for PeptideProphet’s discriminant score to 9.95% for the Philosopher approach.

We hypothesized that this failure to control the FDR may arise in part due to multiplicity in the spectra, that is, due to the presence in the dataset of multiple spectra generated from the same peptide species. For example, Percolator trains its machine learning model using a threefold cross-validation scheme, learning the model parameters on 2/3 of the data at a time and applying the trained model to the remaining 1/3. However, this cross-validation procedure may be violated if one peptide species generates multiple spectra, with some of those spectra going into the training set and some into the test set. In such a situation, the model may overfit. This spectrum multiplicity problem is exacerbated whenever replicate datasets are analyzed jointly.

To test this hypothesis, we repeated the above entrapment analysis, but instead of analyzing all nine ISB18 runs jointly, we analyzed each run separately (Supplementary Figure S1). The resulting FDR violations significantly subside for the case of Percolator, while for PeptideProphet they become worse. Unlike Percolator, PeptideProphet does not use a cross-validation scheme, so a proper accounting for the apparent FDR violation may be due to factors beyond the multiplicity issue.

### 3.3 Revisiting the problem of FDR control in a narrow search

When introducing their open-search software, MSFragger, Kong *et al*. claimed that it uncovered a problem with TDC-based FDR control in narrow search [12]. Specifically, analyzing the 24 spectrum files of the HEK293 dataset [1], they looked for peptides that were discovered in a narrow search with an FDR threshold of 1%, but for which the associated spectrum switched to a different peptide in the open search. In their narrow search results at a 1% FDR threshold, they found 1,139 such “switching” peptides among a total of 101,138 peptides. Assuming that those 1,139 peptides were falsely detected yields a false discovery proportion of 1.13%, which they concluded is too high given the FDR threshold of 1%.

To further support their claim, Kong *et al*. looked for peptides that were identified twice: once in a narrow search without any modifications and once in an open search with a modification mass corresponding to either carbamylation (43.00 Da) or oxidation (15.99 Da). They then collected the experimental spectra responsible for identifying the modified peptides in the open search, and they investigated what peptides were assigned to these spectra in the narrow search results. Because these peptides were identified twice (with and without modifications), we have high confidence that these detections are accurate. Therefore, we expect that if we search these spectra against a database in narrow search mode, while not allowing for the carbamylation or oxidation modifications, then the resulting PSMs should be incorrect. Accordingly, we expect the resulting false PSMs to be equally divided between target and decoy peptides. However, Kong *et al*. report that this set of spectra overwhelmingly matched with more target than decoy peptides: a ratio of 6:1 for carbamylation and of 9:1 for oxidation. Thus, they concluded again that something is wrong with applying TDC to the narrow search. In particular, the abundance of target peptides indicates that the FDR control is liberal.

With its simultaneous consideration of both narrow and open searches, as well as chimeric spectra, CONGA allows us to reevaluate those claims. Specifically, CONGA allows us to factor in peptides that co-generate chimeric spectra, for which both the narrow-search and open-search PSMs can be correct. Indeed, Figure 3 suggests that a considerable portion of narrow-discovered peptides that Kong *et al*. considered as suspect might be due to chimeras: at 1% FDR level, the median percentage of narrow-discovered peptides that CONGA attributes to chimeric spectra is 2.21%, and at 5% this median goes up to 2.83%. In addition, CONGA allows us to account for the fact that some peptides detected in the narrow-search are lost when their identifying spectrum switches to a higher scoring but incorrect match in the open search. Indeed, because the open search considers many more candidate peptides than the narrow search, to declare that a narrow match is incorrect we require that the open search peptide that replaced it is itself discovered.

**Figure 3.**
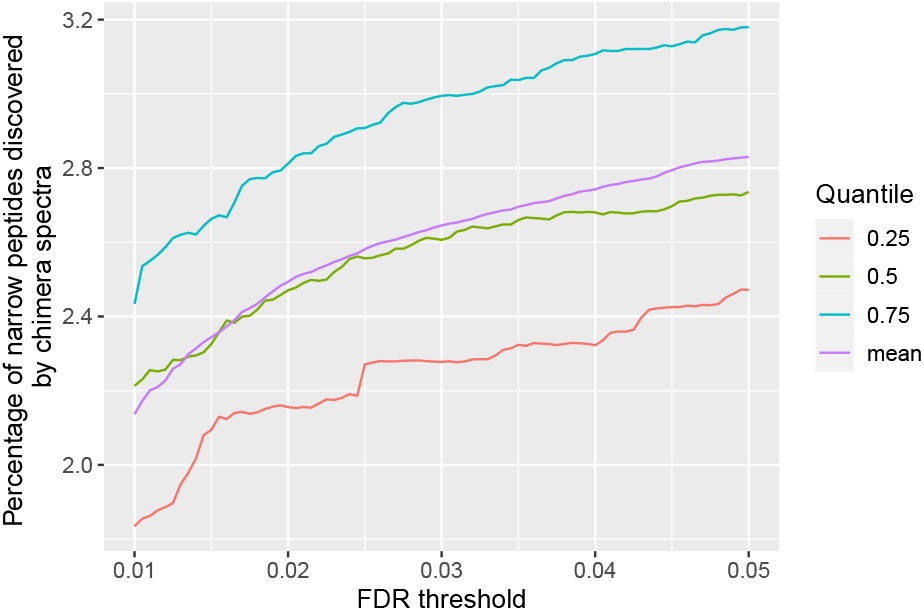
The percentage of narrow-search peptides CONGA discovers through chimeric spectra. The quartiles and mean of the percentage of narrow-search peptides detected by CONGA for which the top-scoring PSM is from a presumed chimeric spectra, plotted as a function of the FDR level. The quartiles and mean are taken over the 24 HEK293 spectrum files, and the percentages are taken with respect to the number of peptides discovered at the same FDR level by CONGA.

Our analysis involves comparing the results of two search procedures—MSFragger in narrow-search mode, followed by TDC, and CONGA—applied to the 24 spectrum files of the HEK293 dataset (details in Section 2.1.3). We proceed in four steps, summarized in Supplementary Table S1. First, we count the number of narrow-search discovered peptides (“narrow peptides”) for each of the 24 spectrum files of the HEK293 dataset at 1% FDR using TDC. The median number of discovered narrow peptides is 10,094. Second, we count the number of the narrow peptides that were lost among CONGA’s discovery list using the same 1% FDR threshold. Such a loss could happen because the identifying narrow PSM was lost due to a higher scoring open PSM—the main effect that Kong *et al*. were concerned with—or because the decoy of the narrow peptide outscored it in the open search, or because the narrow PSM did not score high enough to register as a discovery when CONGA determined its joint list of open and narrow discoveries. Regardless of the reason, the median number of narrow peptides that CONGA fails to detect at 1% FDR is 239, or about 2%. Third, we count the number of lost narrow peptides for which the identifying narrow PSM was swapped for at least one higher-scoring open PSM whose peptide was discovered by CONGA at the same 1% FDR threshold. These peptides correspond to what we consider as dubious narrow discoveries: the identified spectrum from the narrow search was reassigned to another peptide in the open search, and moreover this open peptide is very likely to be present in the sample. The median of these dubious narrow discoveries number is 49. Finally, in step four we compute the percentage of the dubious discoveries among all narrow discoveries. These percentages vary between 0.261% to 0.66% with a median of 0.46%. This is well below the 1% FDR threshold, so we conclude that this analysis does not reveal a problem with the TDC-implemented FDR control of the narrow search of this data.

We also used CONGA to reexamine the second setup that Kong *et al*. analyzed. To do this, we first compiled a list of the spectra that were confidently matched to peptides with mass-modifications coinciding with oxidation and carbamylation (up to *±*0.02 Da). Specifically, the relevant PSM had to be the top match for the spectrum, and the peptide had to be discovered at 1% FDR where, to be more inclusive here, we allowed for any peptide in CONGA’s augmented list of discoveries (described in Supplementary Algorithm 6 of the CONGA manuscript [6]). Following Kong *et al*., we then selected the subset of those spectra for which their matched peptides were also discovered, without any modification, at 1% FDR in the MSFragger analysis. Finally, we found the proportion of these selected spectra whose narrow-search optimal matching peptide is a target one. Assuming that the open search match is the correct one, the narrow search should equally likely match a target or a decoy peptide.

Our results disagree with those of Kong *et al*. They reported that 85.7% of the carbamylated spectra and 90% of the oxidized spectra matched to targets, whereas we observe corresponding values of 55.5% and 53.2%. Some of the observed bias might arise due to chimeric spectra, if some of the open modified peptides share their identifying chimeric spectrum with an unmodified peptide. To account for this possibility, we redid the above analysis after removing those experimental spectra that CONGA identified as chimeric with at least one of the co-generating peptides detected in the narrow search. This brought the target percentages down slightly to 54.3% and 52.3%, respectively. Considering that our chimera analysis is fairly conservative we do not find these biases sufficient to determine that narrow-search based TDC fails to control the FDR.

## 4 Discussion

Open search is an increasingly popular approach to detect peptides that harbor unexpected post-translational modifications. However, applying both open and narrow search to the same data, followed by TDC, often yields fewer discovered peptides for the open search than the narrow one. Only when applying a post-processor such as Percolator or PeptideProphet do we see a fairly consistent improvement using an open search. On the other hand, we demonstrate here that both Percolator and PeptideProphet apparently fail to control the FDR in our entrapment experiments.

Combining the results from the open and narrow searches, our recently developed CONGA offers an alternative approach that theoretically and empirically controls the FDR. Using CONGA we were able to show that the recent claim that open search identifies a problem with narrow search TDC-based FDR control is not sufficiently supported.

## Supplementary information

### Algorithm 1 PSM-and-peptide from [13]

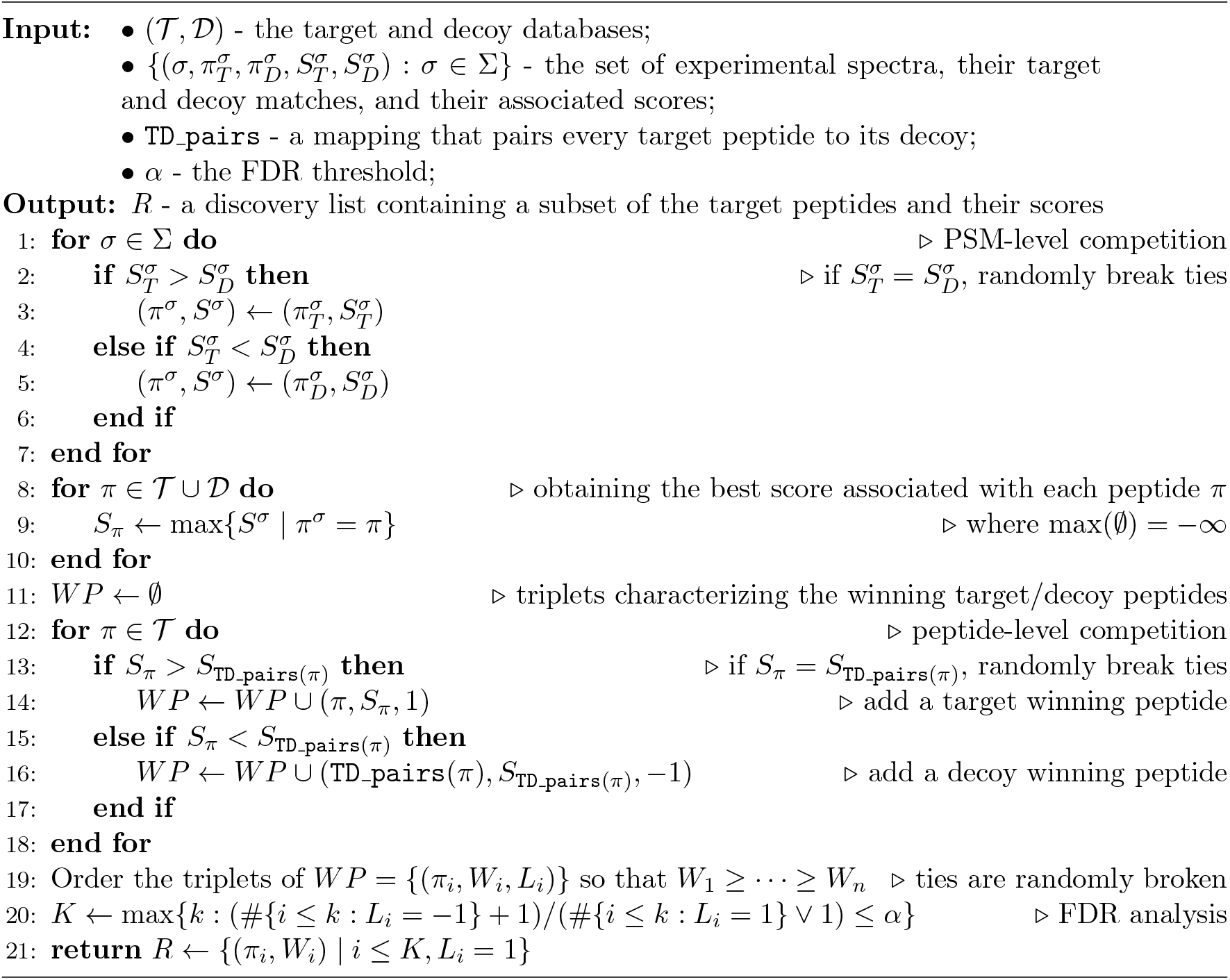

**Table S1:** The number of reassigned peptides discovered by CONGA. The table reports the following information for the 24 spectrum files used in [12]. The first row gives the number of peptides discovered at 1% FDR using TDC applied to the list of top 1 PSMs generated by MSFragger in narrow-search mode. The second row lists the number of peptides from the first row that were not discovered by CONGA at 1% FDR. The third row reports the number of the CONGA-lost peptides from the second row, for which the spectrum that identified them in the narrow search were reassigned to another peptide that was discovered by CONGA. The last row gives the percentage of the the reassigned spectra relative to all narrow discoveries.

| TDC-discovered narrow peptides | Narrow peptides lost in CONGA | Spectra reassigned to CONGA-discovered open peptides | Percentage of reassigned narrow peptides (in %) |
| --- | --- | --- | --- |
| 9456 | 220 | 59 | 0.62 |
| 11116 | 242 | 48 | 0.43 |
| 9056 | 249 | 52 | 0.57 |
| 10879 | 254 | 54 | 0.50 |
| 9986 | 212 | 44 | 0.44 |
| 9603 | 224 | 44 | 0.46 |
| 11162 | 247 | 46 | 0.41 |
| 9181 | 205 | 46 | 0.50 |
| 13221 | 253 | 81 | 0.61 |
| 10203 | 230 | 59 | 0.58 |
| 12930 | 268 | 50 | 0.39 |
| 8675 | 187 | 45 | 0.52 |
| 11265 | 274 | 50 | 0.44 |
| 9195 | 306 | 24 | 0.26 |
| 11518 | 414 | 61 | 0.53 |
| 9765 | 323 | 64 | 0.66 |
| 9524 | 220 | 34 | 0.36 |
| 10610 | 236 | 39 | 0.37 |
| 8709 | 209 | 41 | 0.47 |
| 12143 | 263 | 52 | 0.43 |
| 5012 | 152 | 28 | 0.56 |
| 12688 | 230 | 58 | 0.46 |
| 7292 | 194 | 31 | 0.43 |
| 12530 | 320 | 73 | 0.58 |

**Figure S1:**
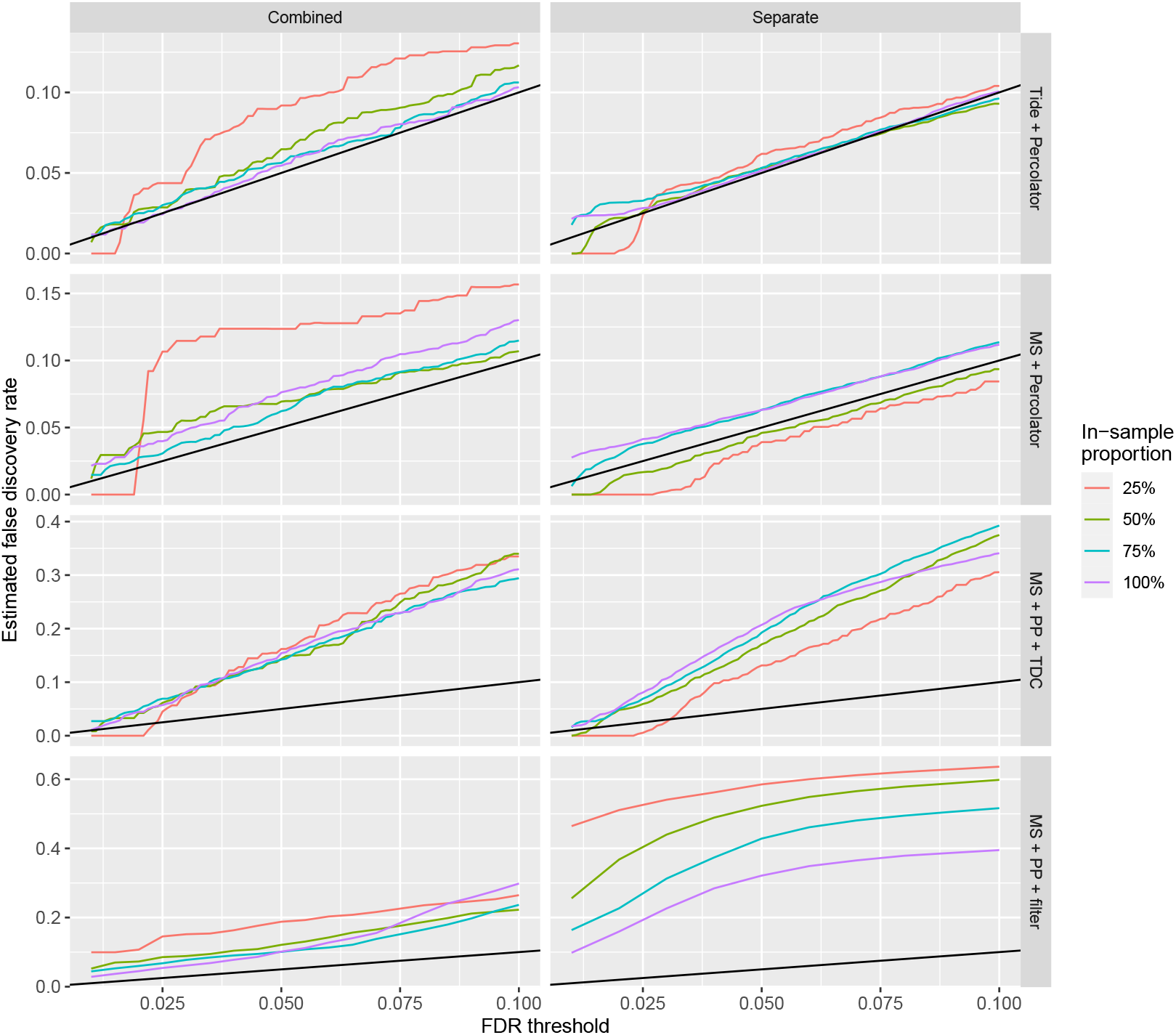
Percolator and PeptideProphet empirical FDR using combined and separate runs. The first column of figure panels is the same as those in Figure 2 search in open-mode. The second column repeats the entrapment analysis, estimating the false discovery proportion for each of the nine ISB18 runs separately and randomly generated decoy database. The estimated FDR is then averaged over each of the 20 randomly generated decoys and nine ISB18 runs. For Percolator, switching from a combined analysis to a separate analysis of the ISB18 runs decreases the estimated FDR and supports the hypothesis that spectra generated by the same peptide species can lead to overfitting. As for PeptideProphet, the estimated FDRs remain well above their respective FDR thresholds. *Note that the FDR scale varies between the rows*.

## Footnotes

1 Supplementary materials are included at the end of this document.

